# A novel citizen science-based wildlife monitoring and management tool for oil palm plantations

**DOI:** 10.1101/2025.01.12.632638

**Authors:** Nunik Maharani, Nardiyono, Claudia Retina Munthe, Priya Swayanuar, Safwanah Ni’Matullah, Syafiie Sueif, Syahmi Zaini, Jatna Supriatna, Mirza Kusrini, Rona Dennis, Bas van Balen, Arco van Strien, Erik Meijaard

**Author notes:** These authors contributed equally to this work (Arco van Strien, Erik Meijaard).

## Abstract

Agricultural expansion is one of the greatest threats to global biodiversity. At the same time, many wildlife species survive or even thrive in agricultural landscapes that retain patches of natural ecosystems. This is especially true for tropical oil palm (Elaeis guineensis, Jacq.) plantations that have both replaced tropical forest and other species-rich ecosystems, but as perennial crops can also function as wildlife habitat, especially if fragments of natural ecosystems are retained. There is an urgent need to understand how to manage and monitor wildlife in these fragmented oil palm landscapes. Still, the lack of large, quantitative datasets on species abundance impedes learning. We piloted a novel citizen science-based biodiversity monitoring system in Austindo Nusantara Jaya’s seven Indonesian oil palm estates, across different biogeographical regions, over a 5-year period. The company-wide monitoring system called PENDAKI, the Indonesian acronym for Care for Biodiversity, is the first of its kind in the palm oil industry. Here we demonstrate that such unstructured and opportunistic data collected mainly by lay people can result in valuable information on temporal and spatial changes in species occupancy. Between September 2019 and June 2024, PENDAKI has resulted in 148,286 wildlife observations, that included 699 reliably identified faunal and 186 floral species, with contributions from 3,950 employee contributors. We estimate species-specific occupancy rates using Bayesian occupancy modeling, ideally suited for opportunistic data where survey effort is unknown. We show that these occupancy data can reliably show temporal and spatial changes in the distribution of iconic wildlife such as orangutans (Pongo pygmaeus). We combined data from reliably identified species with at least 50 records to create the “Living Plantation Index”, an estate-specific annual index of wildlife diversity based on occupancy estimates. We conclude that citizen science-based biodiversity monitoring works remarkably well in oil palm plantations because of the large number of people typically working there. In the process we discovered the emergence of co-benefits such as increasing environmental stewardship awareness across the workforce, raising the profile of the conservation department within the company. We also noted the benefits in terms of suitable data to meet regulatory and voluntary disclosure requirements.

## Introduction

Oil palm plantations, covering 24 million hectares in the species-rich tropics [1], are at the centre of intense debate and under global scrutiny due to their contribution to biodiversity decline [2, 3]. Under pressure, most notably from non-government organizations and consumers, the palm oil industry has been hard pushed to improve its environmental credentials. With the establishment of the Roundtable on Sustainable Palm Oil (RSPO) in 2004, there has been some progress in reducing negative impacts on biodiversity in plantation landscapes [4–6], through biodiversity management requirements such as setting aside areas with High Conservation Values and avoiding deforestation [7]. Determining whether such measures actually benefit the diversity and abundance of wildlife in certified plantations requires measurable indicators and their monitoring [8].

The high granularity of data necessary for effectively monitoring the impacts of plantation management practices on biodiversity necessitates expansive and frequent data collection [9]. This is, however, costly and requires specific expertise that is not generally present within palm oil companies whose conservation staff, are often low in number per estate and usually allocated to numerous other tasks related to environmental management. The uptake of quantitative monitoring of the diversity and abundance of wildlife in and around oil palm plantations has therefore been slow. Most companies, if they do any monitoring, simply draw up annual species lists for monitoring purposes [6]. But adaptive management at the level of the entire estate based only on species presence information is difficult.

Instead, quantitative information at the population level is needed to inform management interventions [10]. In the complex landscapes of oil palm plantations, the need for innovative approaches to monitor and manage biodiversity is urgent [9].

Conventional biodiversity monitoring methods are often limited by logistical challenges and restricted geographical coverage, necessitating innovative solutions to enhance data acquisition and analysis [11]. In this context, citizen science, defined as the participation of non-professional volunteers in scientific research [12], has emerged as a transformative approach for expanding the scope and scale of ecological data collection. Citizen science is a rapidly growing field for increasing both the generation of scientific data and the public participation in understanding and addressing societal challenges. Its application ranges from biological conservation, astronomy, medicine, environmental science, archaeology, and others [13]. The methods are not new though and arguably, and depending on definition, citizen science predates academic science [14].

By harnessing the collective effort of volunteers, in our case oil palm plantation workers, citizen science can generate large datasets that capture the spatial and temporal dynamics of species in these modified landscapes. Such democratization of data collection has proven particularly valuable for biodiversity modelling, which relies on extensive datasets to accurately predict species distributions, phenological changes, and ecosystem responses to environmental stressors [15]. Studies have shown that citizen science projects can yield data of comparable quality to traditional scientific methods, provided that appropriate training and validation protocols are in place [16]. However, integrating citizen science data with sophisticated statistical models poses unique challenges, including ensuring data reliability, and dealing with biases and the variability inherent in data collected by non-experts.

Starting in 2019, our pilot project in the Indonesian palm oil company Austindo Nusantara Jaya (ANJ, further referred to as the “company”) tested whether citizen science-based biodiversity monitoring could be effectively implemented in oil palm plantations, whether this could result in statistically robust data on the occurrence of wildlife, and whether this information could be translated into an adaptive wildlife management cycle. Furthermore, we also wanted to assess the social aspects of implementing a citizen science approach. The potential of citizen science in this context is not just in data collection but also in fostering greater environmental stewardship among participants, promoting a culture of conservation awareness within plantation management practices, and to empower the conservation staff. This is based on the idea that management of wildlife requires that people care, and it is difficult for people to care if they do not ‘see’ wildlife. Collaborative wildlife management in oil palm plantations therefore requires that everyone is involved in the monitoring process. In citizen science, people often contribute without financial reward, and often the research being conducted does not have direct impact on the participants and is led by scientists whom the participants will never meet [17]. This outsourcing of wildlife monitoring to external consultants is common in the palm oil industry. To ensure better internalisation of wildlife monitoring and management in ANJ, developing a sense of ownership over information and pride in participating in wildlife monitoring has been an important objective of this citizen science pilot study. When many oil palm workers care about wildlife, it becomes a lot easier for a company to implement effective wildlife management.

This paper aims to examine the role of citizen science in addressing the biodiversity monitoring and management challenges specific to oil palm plantations. We determine whether it is possible to get palm oil estate workers to collect biodiversity data, and whether such data can then be used for adaptive biodiversity management by the company. We do this by answering the following questions: 1) Was there sufficient participation by palm oil workers? 2) How did we generate commitment within the company to implement this program? 3) Do the data generate enough information to assess population changes over time and is this information used in adaptive wildlife management? 4) Can the company collect, manage and analyze the citizen science data itself? and 5) How do the costs of these citizen science methods compare with more traditional, technology-focused approaches?

## Methods

### Project initiation and location

Leading up to project initiation, there were extensive discussions with senior company management. EM has a longstanding association with the company since 2011. An initial trial of the citizen science approach in the same company several years prior had failed as there was not enough support from field-based staff and estate management. This time, we ensured that senior company managers were fully informed and supportive of the project before we started.

The program was initiated through a one-week in-person training by author RD for 2 conservation managers and 12 conservation and sustainability compliance staff. The principles of citizen science were explained to get staff to understand the approach. Through several training sessions, trainees were asked to develop their own ideas as to how the citizen science project could be implemented within the company, given the specifics of each estate and the capacity of the staff to implement the approach.

They were also asked to name the project, which resulted in the name “PENDAKI”, which is abbreviated Indonesian for Caring for Biodiversity. Through this approach the idea was conveyed that the conservation staff were in charge of conceptualizing and implementing the project, not the outside advisors. A second training session was led by RD in 2023, focusing on standardizing data collection methods. Additional trainings were provided by author AvS regarding data management and statistical analysis in R [18]. Detailed description of statistical methods and modelling assumptions for developing occupancy statistics from the PENDAKI data will be published elsewhere.

The citizen science data collection started in September 2019 in 6 oil palm estates and one sago (*Metroxylon sagu*, Rottb.) concession in Sumatra (ANJA and ANJAS), Belitung (SMM), Kalimantan (KAL) and Southwest Papua (PMP, PPM and ANJAP) (Fig. 1) with a total area of 144,611 ha, of which 59,867 ha (41%) is conservation area (Austindo Nusantara Jaya 2023). The estates are in two different biogeographic regions, Sundaland and Sahul, with very different wildlife species (Fig. 1).

**Fig. 1.**
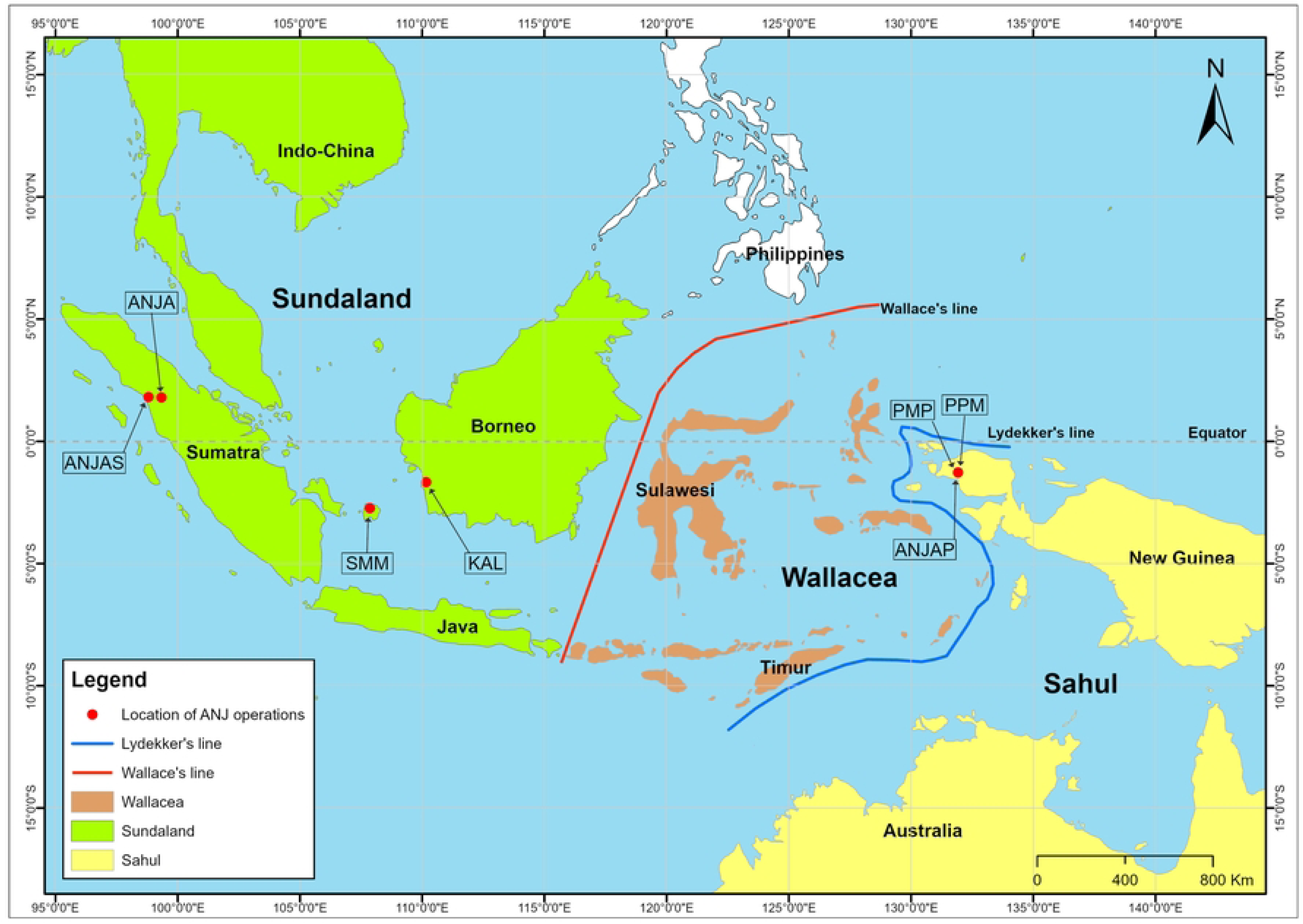
Location map of the 7 Indonesian estates. Abbreviated estate names show where the PENDAKI program was implemented. Also shown are the two biogeographic regions and the Wallace and Lydekker’s lines separating them. Basemap is from [19].

### Data collection

Participation in PENDAKI was possible for any company staff, contractors, or external visitors who volunteered to report wildlife observations to the company’s biodiversity teams in the company’s estates. This included the company’s core conservation staff, other staff whose work was not associated with biodiversity monitoring and management (e.g., harvesters, mill workers, drivers, security guards, cooks, estate managers), and visitors (e.g., journalists, biodiversity consultants, government staff). Any of these people could report any wildlife observations by recording it on a paper form, or directly reporting it to the conservation units in each estate (either through in-person meetings or mobile phone communication). In 2023, a custom-built smartphone application, PENDAKI Champion, was rolled out across the 6 oil palm estates to increase data collection and to link each wildlife observation to a precise spatial coordinate. The application has 25 target species for each estate, and the idea is that PENDAKI Champion collect data on these target species whenever these are observed. This improves the statistical analysis of occupancy.

Participation in PENDAKI was voluntary and there was no monetary reward for contributing information. However, senior ANJ management instructed all estate managers to help implement the PENDAKI program in their estates and linked its implementation to Key Performance Indicators for specific staff.

Furthermore, the company implemented a rewards program that gave small tokens of appreciation (t-shirts, caps, raincoats) for the most interesting wildlife observations.

### Data management, validation, and cleaning

The PENDAKI program was initially paper based with each individual wildlife observation being recorded on a form, with all the relevant details of the observation (date and time, location, species name, behaviour of species) and a photo, if available. This information was then entered monthly into an Excel spreadsheet and each estate would share these sheets with the head office in Jakarta, where the data would be compiled into the master database. The PENDAKI smartphone application introduced in 2023 was initially rolled out by a selected group of 10 of the most active data collectors per estate (called PENDAKI Champion). The paper-based and application-based system ran in parallel with all non-app data being compiled and uploaded to the PENDAKI Dashboard, whilst the app-based data was automatically uploaded to the PENDAKI master database.

PENDAKI species observation were noted in reference to the numbered block system that is in use in most oil palm plantations. These oil palm blocks cover mostly 50 ha (2 km x 0.25 km), although some blocks are smaller, depending on local topography. The protected High Conservation Value (HCV) forest areas were in most cases much larger than individual oil palm blocks, and were referred to in the reporting by their specific name (e.g., “HCV 657 ha”). This grid system provided the initial spatial reference for the species distribution modelling and thus determined the maximal resolution of the analysis. In 2024, all locations at the estate level were standardized so that each wildlife observation was linked to a particular polygon that was ecologically broadly homogenous. The ecological characteristics of these polygons were either 1) natural forest; 2) oil palm; 3) waterways, ponds and palm oil mill effluent (POME) ponds (particularly for birds such as waders and egrets); and 4) infrastructure (offices, factory, housing). When PENDAKI data collectors used the PENDAKI application, each wildlife observation was automatically linked to geospatial coordinates.

On an annual basis, external species experts validated the PENDAKI records and determined the likelihood of the reported wildlife sightings being correct using the following categories (Likely Correct; Unlikely but Possibly Correct; Probably Incorrect; Correct ID but Wrong Species Name; Very Difficult to Identify; and Not Identified at Species Level; for definitions see S2 Table). The reviews used supporting photographs, and knowledge of species distribution and most recent taxonomy to determine the accuracy of records. In the statistical analyses (see below), we only used the Likely Correct species records. Following each data review, a report was provided to the company with key species identification issues, for example, species that were consistently misidentified (e.g., Chinese Egret *Egretta eulophotes*, instead of Little Egret *Egretta garzetta*).

### Data analysis and occupancy modelling

We used Bayesian occupancy models that are ideally suited for the opportunistic, non-structured wildlife surveys to determine spatial and temporal changes for species [20, 21]. We will discuss the technical details and model assumptions elsewhere. We only used data from 2020-2024 because in 2019 PENDAKI contributors did not record several common species. Occupancy models use species-specific detection and non-detection data and provide estimates of the percentage of occupied sites (occupancy), here 50 ha blocks, per year. These models account for the imperfect detection of species. By adjusting for detection bias, they can simultaneously address observation and reporting biases [21]. For demonstration purposes, we determine spatial and temporal occupancy (or distribution) trends of orangutan (*Pongo pygmaeus*), a species normally only associated with forest habitats, but increasingly understood to also survive in heterogenous landscape of forest and agriculture, such as oil palm [22, 23]. We also show the example of the White-breasted Waterhen (*Amaurornis phoenicurus*), a common bird species in the planted oil palm areas.

### Living Plantation Index

For each estate we combined occupancy data for individual species into a compound metric, which we called the Living Plantation Index (LPI), in reference to the World Wildlife Fund’s Living Planet Index [24], although the two indices are mathematically not the same [see 25]. The LPI is based on the average species richness per grid cell, where species richness per grid cell is not the observed number of species but the summed occupancy estimates of the selected species. To calculate the LPI, we only included the annual occupancy means for reliably identified species with a minimum of 50 records over the 5 years of data collection, and at least 5 observations within each year. We weighted the importance of these species in the index as follows: 1) IUCN Red List status (IUCN 2024) – Critically Endangered = 10, Endangered = 8, Vulnerable = 6, Near-Threatened = 4, Least Concern = 2, others = 0; 2) protected status in Indonesia – protected = 10, not protected = 0; 3) CITES listing – List I = 5, List II = 3, not listed = 0; and 4) range – endemic = 5, native but non-endemic = 2, non-native = -2, and migratory = 2. We plotted the resulting index with the 95% confidence intervals resulting from the analyses. To show the temporal trend in the index, we used 2020 as the baseline year for which we set the index value at 100.

### Interview surveys

To better understand the uptake of PENDAKI and how this was perceived by its users and company management, RD conducted one-on-one interviews to gain insights into the motivation underlying participation in the program. The short semi-structured interviews were conducted in June 2022 with 39 staff across the six ANJ oil palm subsidiaries, as well as with head office staff (Supplementary Materials). Of the 39 interviewees, 30 respondents were male, and 9 females. Interviewees were pre-selected by ANJ from each of the 6 subsidiaries and head office in Jakarta, ensuring 5 or 6 from each subsidiary. The interview comprised 13 questions which took about 20 minutes to run through (S1 Table).

A follow up interview survey was conducted in May 2023 by RD and EM to assess the functioning of the PENDAKI Champion smartphone app as well as the original PENDAKI system. The interviews were conducted in focus group discussions in ANJA, ANJAS, PPM, and PMP with the conservation managers and staff present. The total number of participants was 30 app users who were all male, and 12 PENDAKI original users, 4 of whom were female. The duration of the discussion was approximately 3 hours at each of the 4 estates with the opportunity for all to share experiences of using the system including opportunities for improvements both technically and in terms of implementation.

### Ethics statement

This study uses wildlife observation data collected by volunteer staff of PT Austindo Nusantara Jaya Tbk (ANJ). PENDAKI is a program under ANJ’s Responsible Development initiative. Each Responsible Development program has a Project Card listing all participating employees, signed by the respective Project Manager and Director in Charge. In the participating estates, all company staff are informed on a weekly basis about the program and the opportunity to participate. The purpose of the program, which includes identifying present wildlife and analysing the data, is clearly communicated. In Indonesia, the approval of an ethics committee is not required for obtaining and analysing citizen science data, and therefore, it was not obtained. We have received all the necessary permissions required in Indonesia.

## Results

### Overall results

Between September 2019 and June 2024, a total of 148,286 PENDAKI wildlife observations were recorded for the seven estates (Table 1). There is some overlap in names of PENDAKI observers, but we estimate that at least 3,950 individuals contributed wildlife observations since the program’s inception, although not all are regular contributors or still employed by the company. Following review of the observations, 69% were considered taxonomically Likely Correct, leaving a total of 102,626 wildlife observations of 699 reliably identified faunal and 186 floral species for statistical analysis. In addition to species reviewed as “Likely Correct” the species that were “Not Identified at Species Level” (9.2% of total species) and “Correct ID but wrong species name” (3.8% of total species) could also be considered correct identification but with insufficient taxonomic precision. In addition, 6.6% of species records were “Unlikely but Possibly Correct” and could be genuine range extensions, while 2% of the species records concerned species that specialists considered to be very difficult to identify in the field. Some 20 species records proved to be verified range extensions; these occurred especially in Papua and on Belitung which remain relatively poorly surveyed. This leaves 12.2% of species that were incorrectly identified.

**Table 1.**
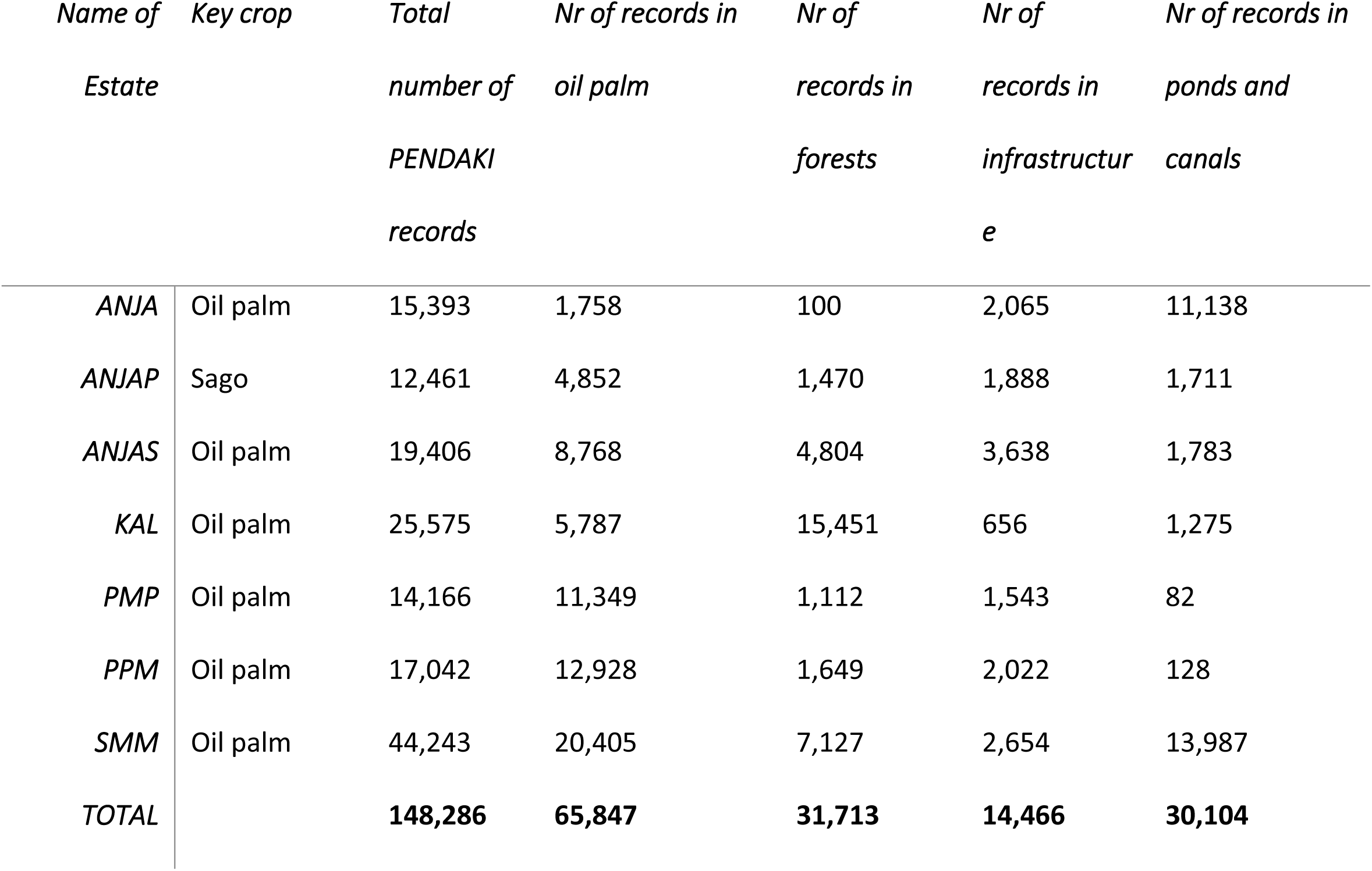
Number of PENDAKI records across company estates. Records are differentiated by location, and were collected between September 2019 and June 2024.

Taxonomically, most of the reliable records concerned sightings of birds and mammals, with much lower numbers for reptiles, amphibians, fish, insects and other species groups (S3 Table). The three most recorded bird species were White-throated Kingfisher (*Halcyon smyrnensis*, Sumatra and Kalimantan), Black-winged Kite (*Elanus caeruleus*, Sumatra and Kalimantan), and Black-capped Lory (*Lorius lory*, Papua). The three most recorded mammal species were Long-tailed Macaque (*Macaca fascicularis*, Sumatra, Belitung and Kalimantan), Large Treeshrew *(Tupaia tana*, Sumatra and Kalimantan), and Pig-tailed Macaque (*Macaca nemestrina*, Sumatra and Kalimantan). For reptiles the three most recorded species are Common Water Monitor (V*aranus salvator*, Sumatra, Belitung and Kalimantan), Reticulated Python (*Malayopython reticulatus*, Sumatra, Belitung and Kalimantan), and Equatorial Spitting Cobra (*Naja sumatrana*, Sumatra, Belitung and Kalimantan).

Most records (44.4%) were collected in oil palm areas, followed by forest (21.4%), ponds and waterways (20.3%) and infrastructure (9.8%), with the remainder coming from sago areas, or unknown or unidentified locations (Table 1). The high percentage of records from the oil palm areas is noteworthy because this is not normally where biodiversity surveys are conducted by biodiversity specialists. It indicates the considerable buy-in from non-biodiversity related workers into the PENDAKI system and supports the central premise of citizen science that all records are important, even when they are considered as common species.

### Examples of occupancy analyses

#### Mapping species occupancy to understand landscape use

We provide examples of how the PENDAKI can be used to provide wildlife-related insights that can be used in adaptive conservation management. For example, while orangutans are known to survive in fragmented forest landscapes, we know little about the spatial use of these landscapes and how plantation management could be adapted to maximize population viability. Between September 2019 and June 2024, a total of 1,028 orangutan sightings were recorded by the PENDAKI observers, of which 829 were in forest areas and 133 in oil palm blocks. Fig. 2 shows the variation between years in mean orangutan occupancy for all survey blocks in the KAL estate in West Kalimantan. Orangutan populations in the KAL estate have remained stable in the past 5 years, possibly showing an increase after 2020. In 2019, extensive wildfires adjacent to the estate destroyed an area of community forest and its resident orangutans likely moved into the KAL estate, resulting in an increase in orangutan densities in KAL’s protected forest areas (IAR Indonesia 2019). While orangutans are mostly concentrated in the protected forest part of the KAL estate, another species, the White-breasted Waterhen (*Amaurornis phoenicurus*) is a common species on the planted oil palm areas, as is shown in a much higher average occupancy (around 0.35) compared to orangutans (around 0.1) (Fig. 2).

**Fig. 2.**
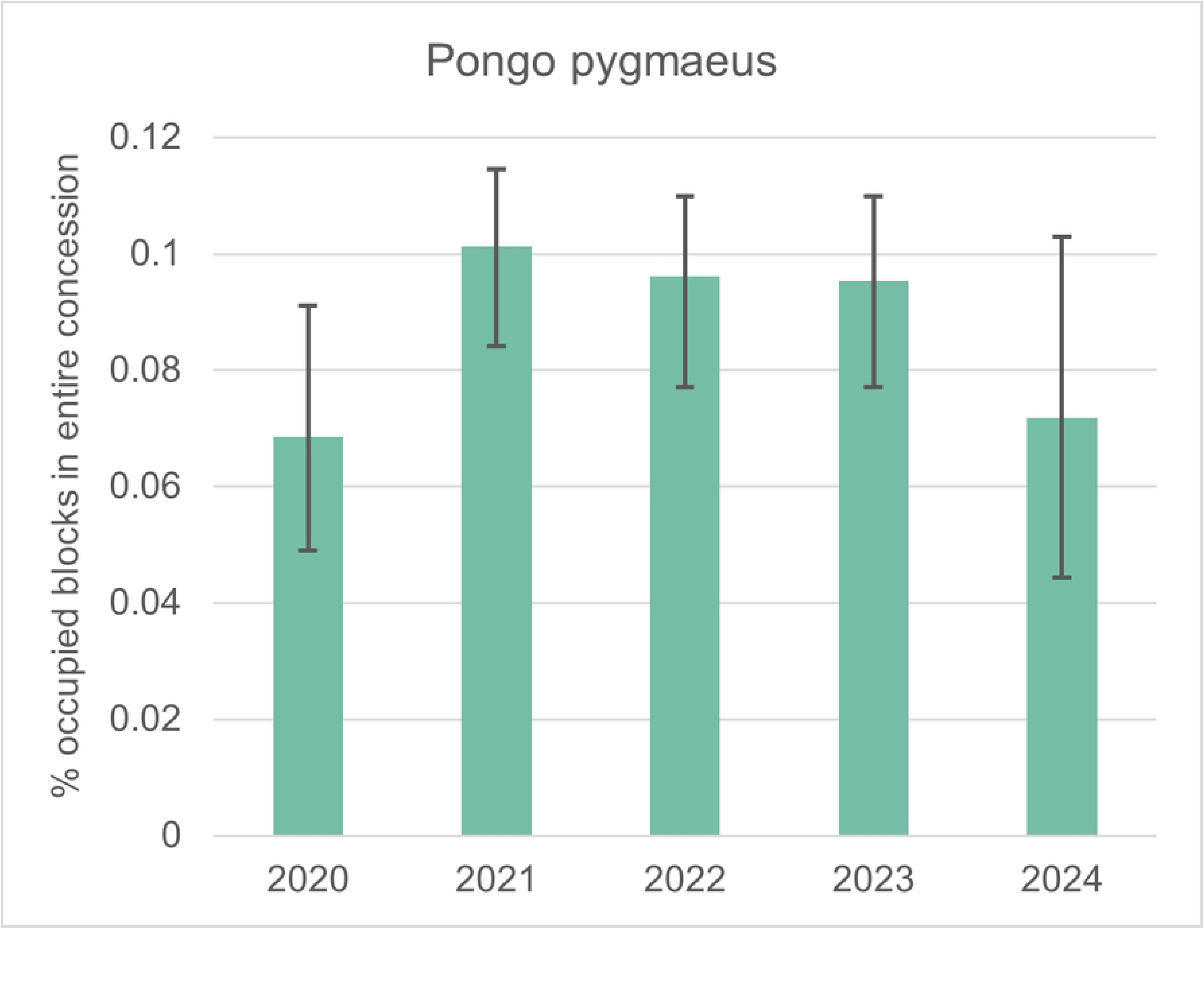
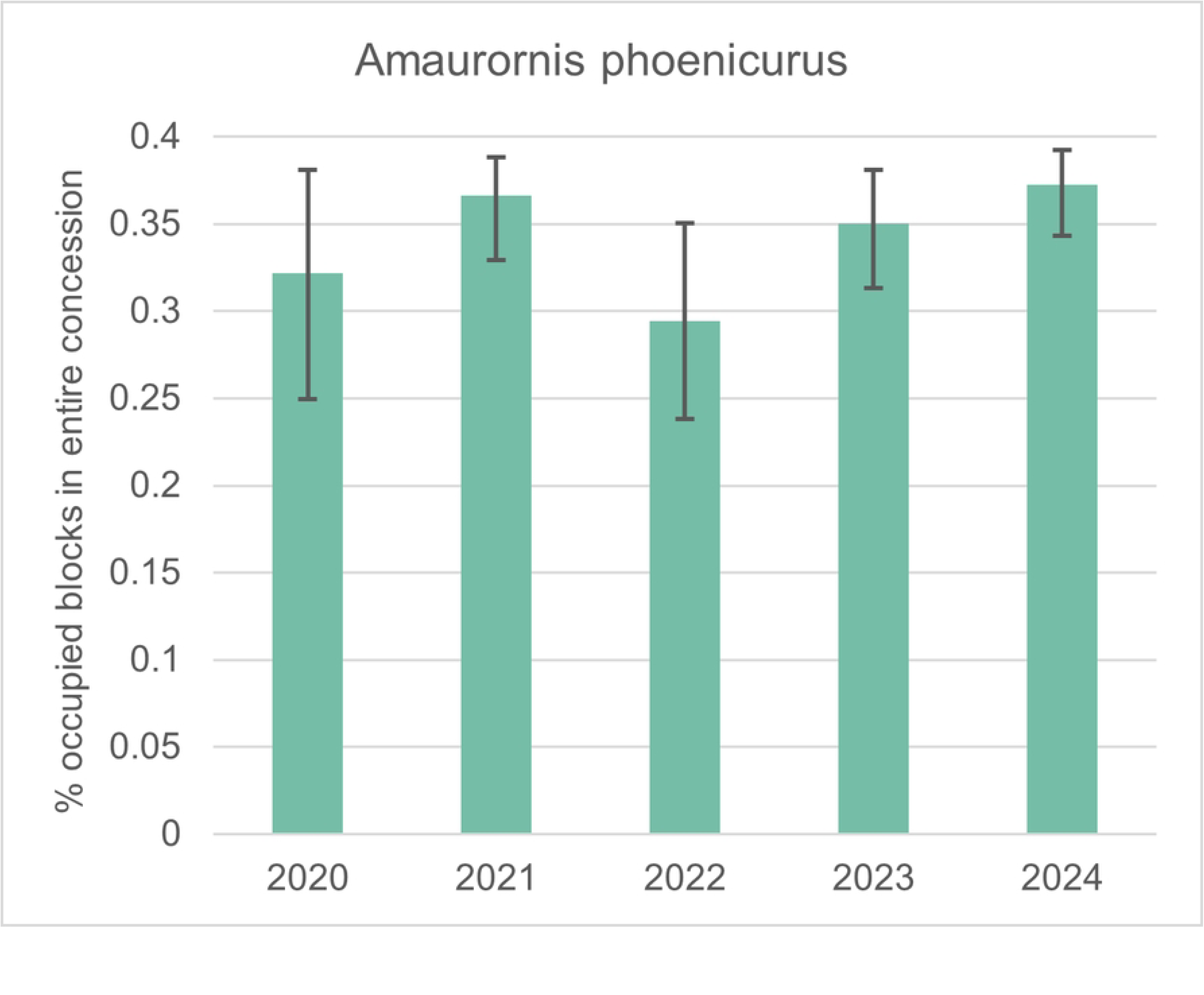
Mean percentage of blocks occupied by two example species. Fig. 2A shows Orangutan occupancy and Figure 2B shows the White-breasted Waterhen occupancy for the KAL estate in Ketapang, West Kalimantan.

Fig. 3 shows the temporal and spatial variation in orangutan occupancy for different survey blocks (forest and non-forest) in ANJ’s oil palm plantation in Kalimantan. The two large forest set-asides (marked A and B in Fig. 3) are areas with permanent breeding populations. Female orangutans tend to be reluctant to leave forest areas, unlike male orangutans [23, 26], so we assume that other forest blocks are only used by male orangutans in certain years. These males seem to prefer moving between forest areas using particular oil palm blocks and avoid others, but we do not know what underlies this preference. A natural forest regrowth corridor established in 2015 to facilitate orangutan dispersal does not seem to be used at all by orangutans, at least according to the data currently available in the PENDAKI dataset. Although much more needs to be learned about orangutans in fragmented landscapes, this kind of information helps a lot in understanding their ecological needs, and informs longer term population viability considerations. For example, if females do not leave forest blocks even if other suitable forest sites are in the landscape, this would reduce the viability of the meta-population.

**Fig. 3.**
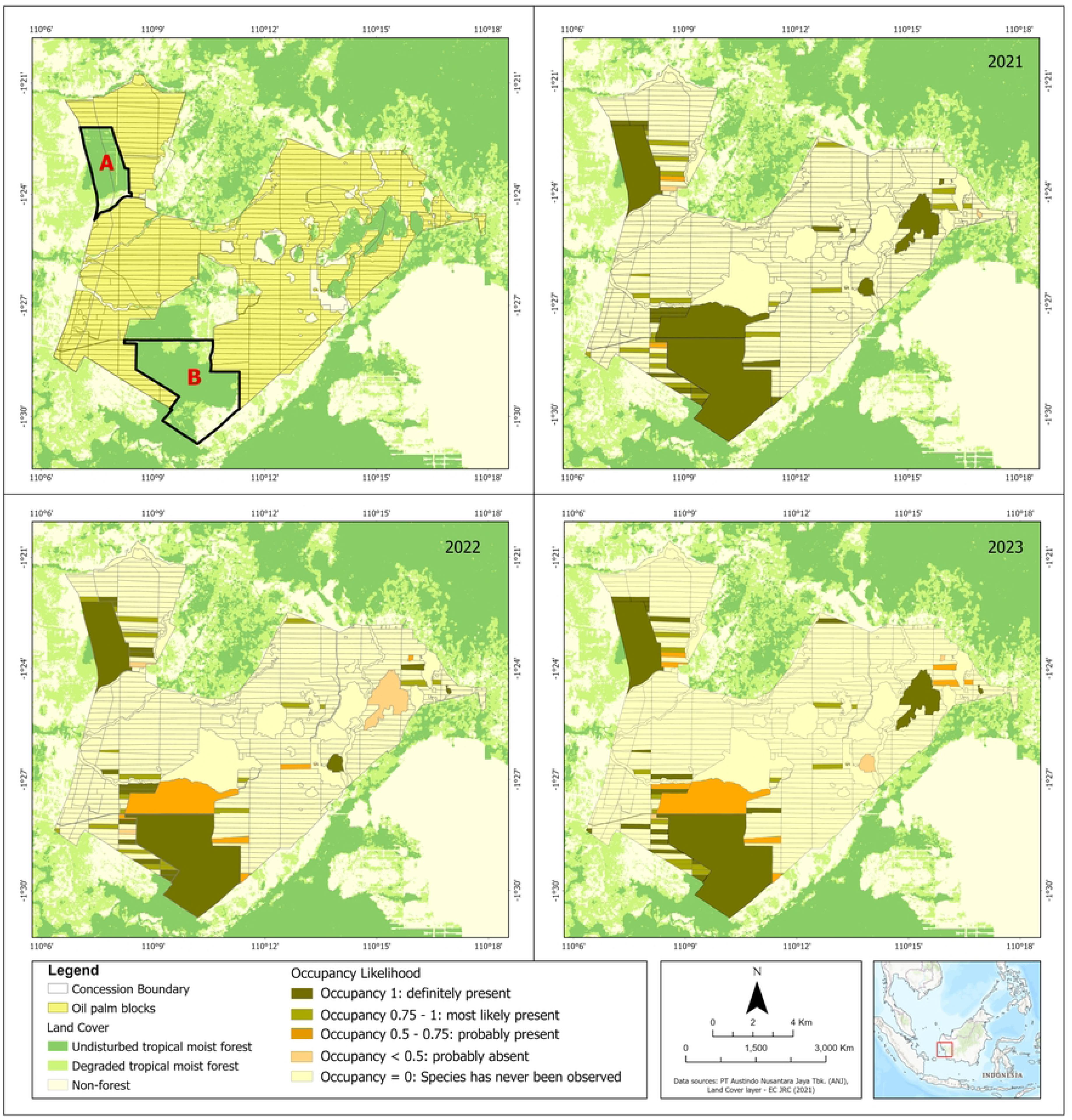
An example of the use of occupancy statistics derived from PENDAKI. The map shows occupancy values for orangutans in the PT KAL concession, in Ketapang, West Kalimantan, Indonesia. Maps are for 2021, 2022, and 2023. The letters “A” and “B” indicate breeding populations of orangutans, i.e., females are present, whereas other areas only have roaming male orangutans.

#### Determining diversity change over time through the Living Plantation Index

Whereas in the section above we described single species occupancy analyses, it is also possible to combine these data to develop weighted species richness indices. We used the occupancy-based diversity data for all survey blocks (forest and non-forest) combined to create the Living Plantation Index that uses the mean occupancy for species per year and a weighting system based on the threat levels to the species, protection status and range (see Methods). Compared to the baseline year of 2020 there seems to have been an increase in occupancy and richness in KAL (Fig. 4).

**Fig. 4.**
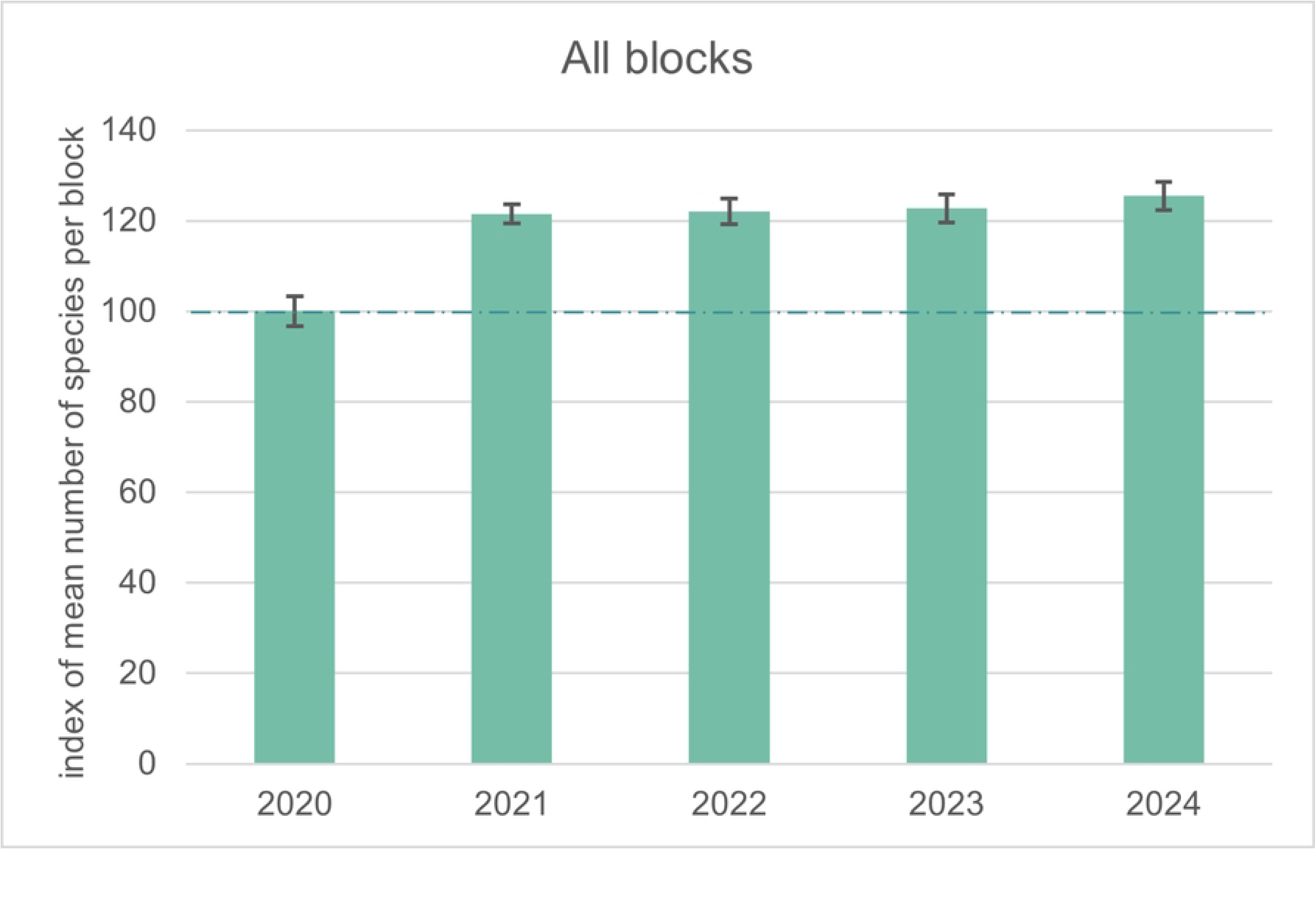
Living Plantation Index for KAL from 2020 to 2024. The LPI was calibrated against an index value of 100 for 2020. Analyses based on the data of 20 species with > 50 records in total and at least 5 each year. Data for 2024 only include until June 2024. Confidence intervals are shown.

Conducting occupancy analysis is important because naïve population or diversity estimates that do not consider survey effort and detection likelihood give a biased picture. The power of occupancy analysis is that the method compensates for variation in survey effort and knowledge of species. Fig. 5 shows the difference in target species richness using modelled occupancy and naïve estimates. For this, we only used species for which there have been a significant number of observations (> 50 in total and at least 5 in each year), resulting in the inclusion of the diversity index of 20 species. Naïve estimates are much lower than modelled estimates in both forest and non-forest sites.

**Fig. 5.**
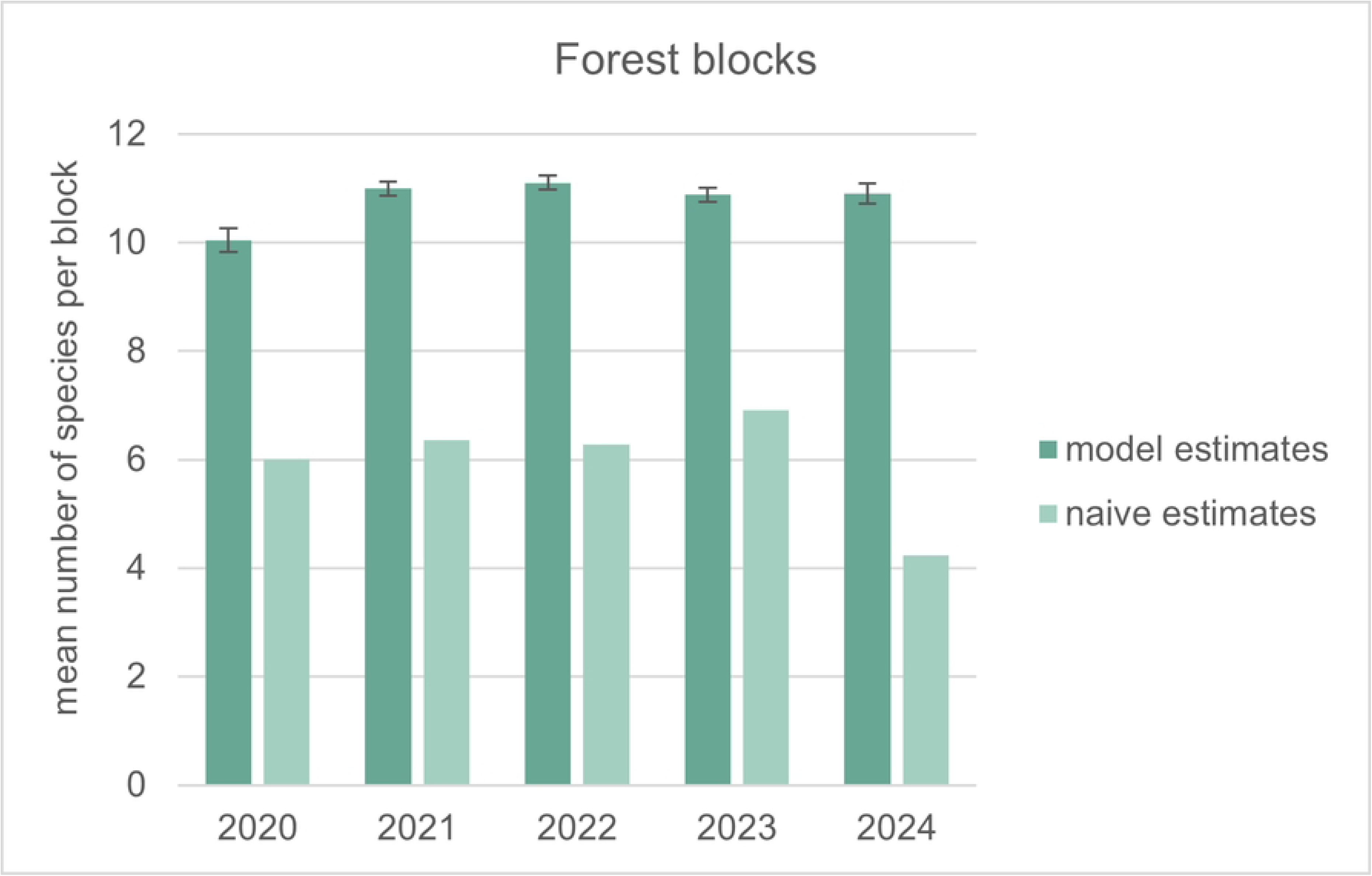
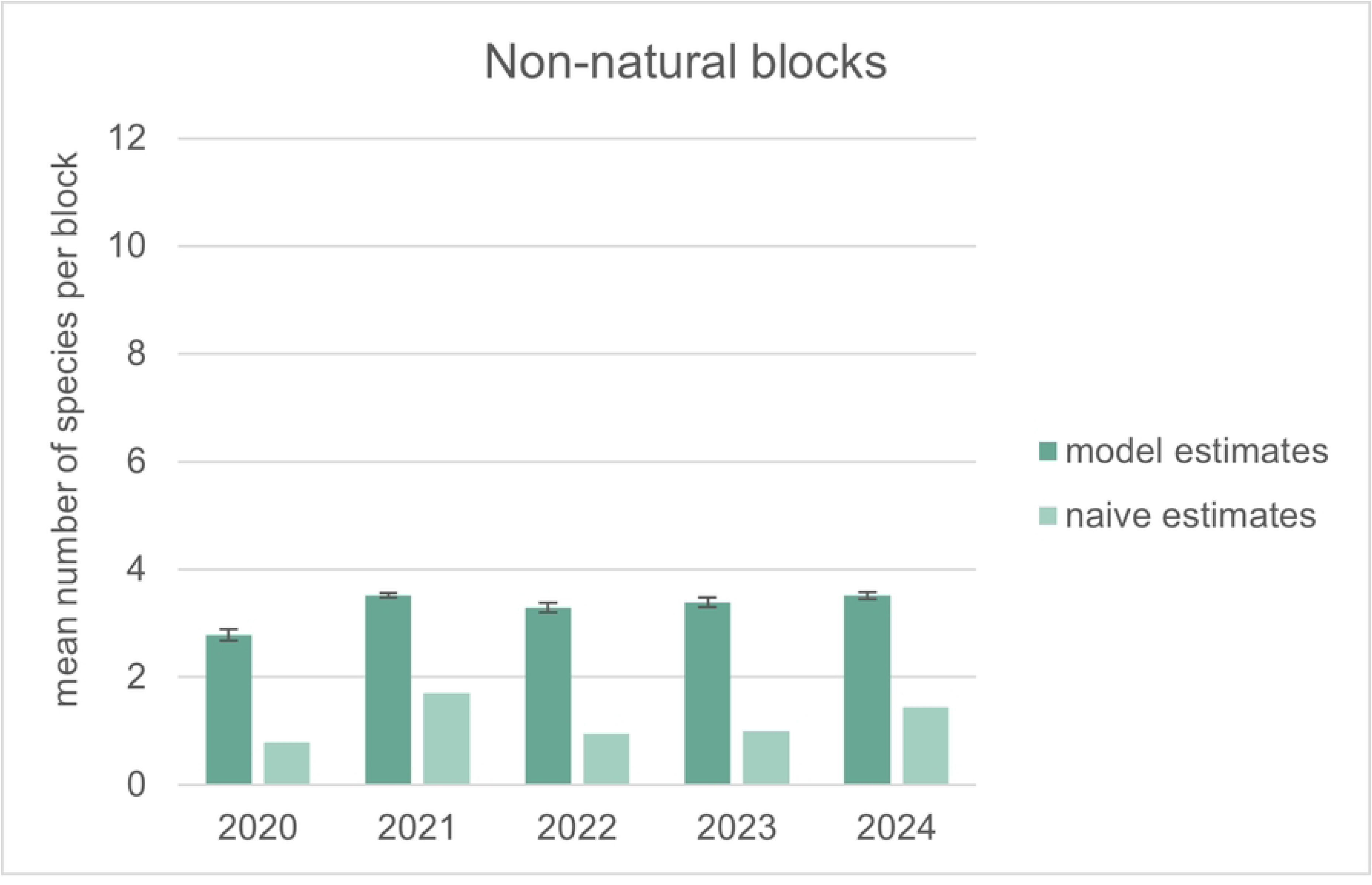
Number of species per survey block. Fig. 5A shows forest blocks and Fig. 5B shows non-natural blocks in one plantation, PT KAL in West Kalimantan, calculated either by occupancy model-based or naïve methods. Analyses based on the data of 20 species with > 50 records. Data for 2024 only include until June 2024. Confidence intervals are shown.

### Interview surveys

The overarching conclusion from the 2022 interviews was that PENDAKI is a highly regarded and popular program with very good buy-in across the company at all levels interviewed from subsidiaries to head office. Respondents feel pride in working for a company that places a high priority on operating sustainably with a clear emphasis on caring for biodiversity and the environment. Respondents considered PENDAKI as a method for recording species data (41% of respondents) and identifying species (28% of respondents), with 36% of respondents mentioning the phrase “citizen science” or describing the program as one which involves everyone in monitoring species.

The level of interest expressed by respondents in seeing wildlife in their everyday job environment was very high. Most respondents expressed an interest in seeing wildlife prior to participating in PENDAKI, but 87% stated that since participating in PENDAKI their interest had significantly increased.

Respondents stated that through PENDAKI they had learned the species names, they knew which species were threatened or protected, and they had a much better understanding of the importance of protecting species and the environment.

The interviews assessed the challenges or difficulties encountered in making an observation. It was found that 46% of respondents experienced challenges with taking a good enough photo of the animal, and 38% of respondents reported difficulties identifying particular species. A PENDAKI WhatsApp Group is in operation at all subsidiaries and was frequently mentioned by the respondents as a means of getting support from the Conservation Team in species identification. The Conservation Team at each subsidiary plays a crucial role in enabling and supporting PENDAKI. Respondents frequently mentioned the quick response on species identification from the Conservation Team. The positive feedback and encouragement from the Conservation Team was evident at all subsidiaries.

The interview survey in June 2024 found that many of the findings from the 2022 were still valid. For long-term users of the original PENDAKI system, there seemed to be more confidence in species identification. It was also noted that motivation to participate in the original PENDAKI system varied across the subsidiaries for various reasons, such as low literacy rates in PPM and PMP. The role of the Conservation Team in motivating workers to participate continues to be important. The reward system was an active topic of discussion with ideas for improving the system particularly for PENDAKI Champion users, but generally participants were satisfied with the reward system.

The discussions yielded insights into the functioning and implementation of the smartphone app. Generally, users were satisfied but still had challenges with identification of some of the 25 target species (PENDAKI Champion), some issues with location connectivity and requested the ability to upload video and audio. It was noted during the June 2024 estate visits that Conservation Staff had increased their species identification skills, and some staff were becoming more knowledgeable of bird calls, and using species scientific names more regularly than previously observed by RD and EM.

### Implementation costs

PENDAKI is low cost because it largely depends on information that is voluntarily collected by company staff. The main costs associated with the program were upfront costs for technical and statistical support during the pilot phase, and the development of the PENDAKI application (see below) (Table 2). ANJ is now meeting all its species reporting and disclosure requirements (government, RSPO, Indonesia Sustainable Palm Oil, Global Reporting Initiative, SPOTT, and other ESG reporting standards) with PENDAKI data. The annual running costs, in addition to basic conservation staff salaries, are primarily associated with rewards and internal and external reporting on PENDAKI. The annual running costs additional to basic conservation staff salaries add up to ca. USD 46,000 in the initial 5-year pilot phase and USD 20,000 in the longer run. This translates into biodiversity monitoring costs of USD 0.32/ha (start-up) or USD 0.14/ha (long-term), when calculated over ANJ’s entire landbank.

**Table 2.**
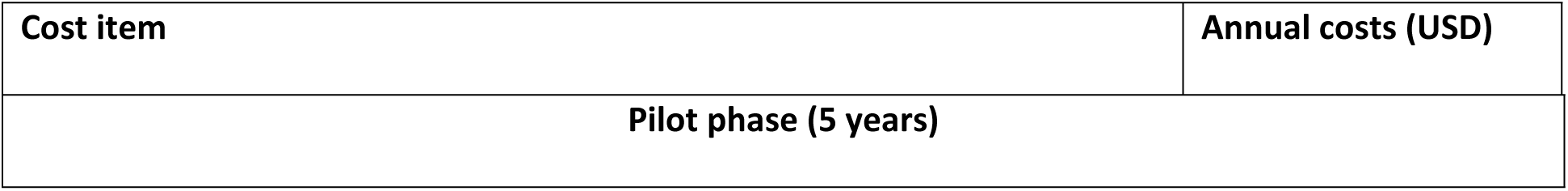
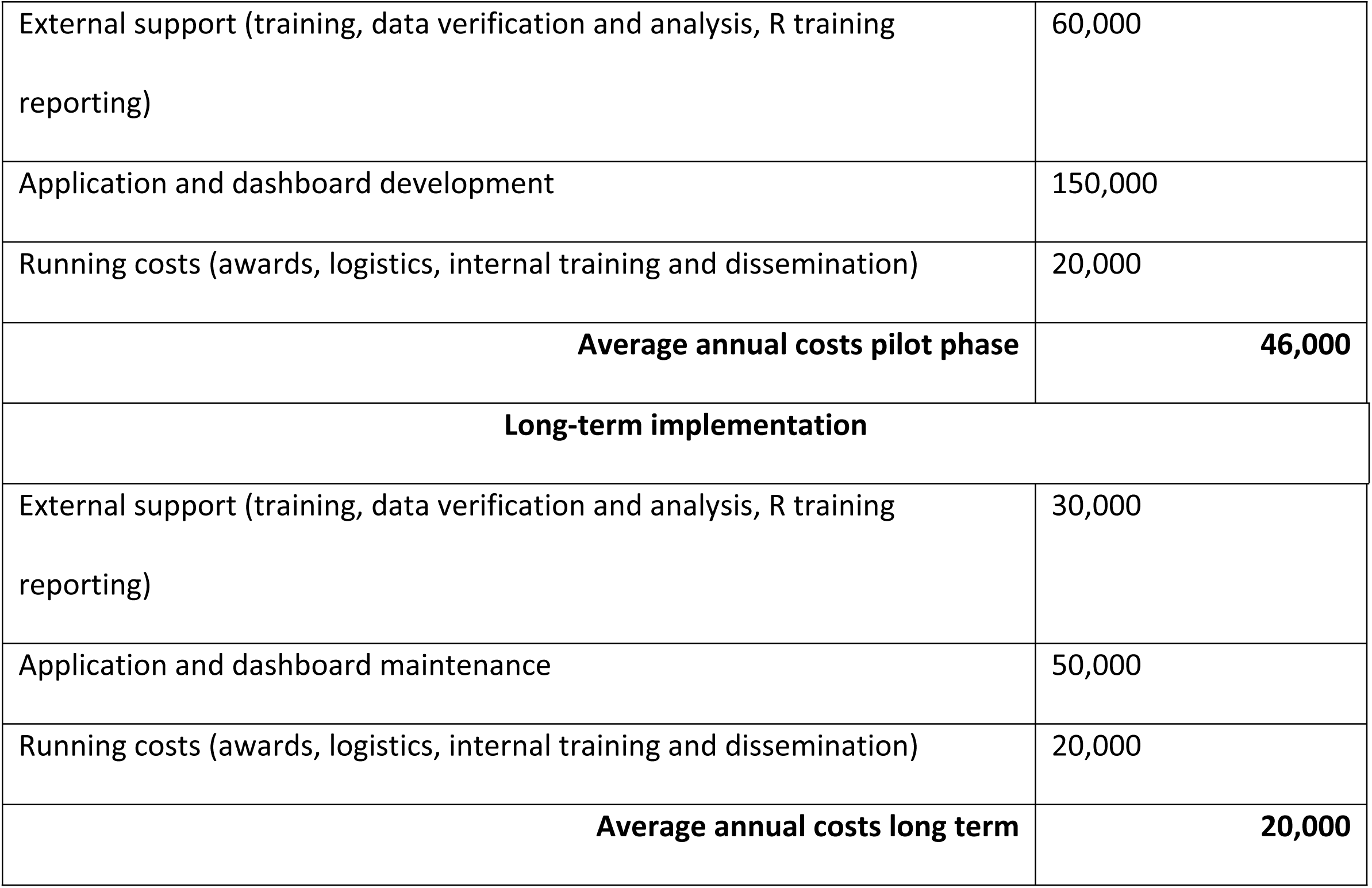
Estimated annual running costs of PENDAKI during the initial 5-years pilot and the long-term implementation phase.

## Discussion

### Participation and buy-in

Unique to PENDAKI, compared to public participation in citizen science, is that the imparted benefits are not limited to data collection and benefits felt by the observers, but also the benefits extend to the company. Data is the predominant focus of much of the published citizen science literature with much less analysis about the factors that influence and sustain people’s participation [17].

Key to the success of any citizen science system is the participation of observers. As PENDAKI evolved it became very clear that the system is as much about people as it is about data [27]. Similar to others [See 28, 29], our first set of interviews in 2022 confirmed that participants feel that they are acquiring new knowledge about species and becoming more aware of the biodiversity values in the area where they work. Interestingly, we encountered participants who stated that they had always enjoyed seeing wildlife around them but didn’t know the species name or anything about the species, and no one had ever asked their opinion about wildlife. Participating in PENDAKI has empowered individuals to express their values regarding the biodiversity surrounding them. It serves as an example of an approach that overcomes the siloed structures often found in corporate environments. PENDAKI emphasizes that biodiversity monitoring is a shared responsibility. The contributions of regular PENDAKI observers play a crucial role in enhancing biodiversity monitoring and, ultimately, its management.

Beyond the observers we found that the PENDAKI system appeared to empower the conservation staff at the estate level. PENDAKI provided a continuous monitoring system for species across the entire estate, a system which was not previously in place as effort was mainly focused on the conservation set-asides. PENDAKI has enabled the conservation staff to improve their species identification skills, readily identify new species for the estate, acted as an early warning of any issues related to species, and has increased engagement with the workforce.

### Corporate commitment

One of the keys to success of PENDAKI was the full support from the company’s senior management, and their requirement for key people (conservation and estate managers) to support PENDAKI’s implementation. This management push got the program across the initial implementation threshold until the citizens science approach became the standard for biodiversity monitoring across the company. At the corporate management level, the profile of PENDAKI has risen exponentially, with increased buy-in and recognition by the highest levels in the organization as the system delivers multiple benefits to the company. PENDAKI has succeeded in raising the profile of biodiversity across the company and has clearly contributed to raising the environmental performance and credentials of the company through recognition by peers, government and national and international awards. Maintaining the engagement and support of both participants and management is crucial for ensuring the long-term sustainability and impact of the PENDAKI system.

### The value of citizen science in adaptive wildlife management

One of the greatest challenges in wildlife conservation is the monitoring of impact from conservation investments on wildlife. Reliably surveying wildlife populations requires large data volumes, which are expensive, time consuming to collect and difficult to interpret. The call for increasing the evidence base for conservation [30] is undermined by these inherent delays (it can take years before sufficient data are accumulated) and high costs of this evidence [31]. Furthermore, the scientific expertise that is required to analyse wildlife data often creates a distance between practitioners who are implementing conservation and data experts who analyse them [32]. This is especially the case in conservation programs implemented by rural communities or companies that have limited scientific wildlife monitoring expertise. Our description of the PENDAKI citizen science program shows that this approach can effectively address the above constraints. PENDAKI has proven to be of potential value for generating statistically useful information about spatial and temporal changes in species occupancy through a system that is low cost and simple to implement. Because observers are not taxonomic specialists, identification errors occur. These errors were identified through verification and are limited to 12% of the observations. Arguably this error rate should not affect project goals if the primary conservation target species (e.g., orangutans or gibbons) are always accurately identified. Citizen science can therefore fulfil particular but not all monitoring objectives. This is reflected in a recent review [33] that found that of the 365 indicators of the Kunming–Montreal Global Biodiversity Framework (GBF) monitoring framework, 110 (30%) can involve citizen science-based monitoring programs and 185 (51%) could benefit from citizen involvement in data collection, while 180 (49%) require scientists and governmental statistical organizations. Recognizing these constraints, the PENDAKI system now provides ANJ with real-time information about readily identifiable wildlife species in seven estates, some of which are very remote, and allows real-time monitoring of these species and management of their threats.

While occupancy data are generally good predictors for species abundance [34], the two are not the same. Occupancy data provide information about spatial use of landscapes and how this varies over time, but it does not allow estimation of species density or population size. Relative changes in occupancy, however, is exactly the kind of information that a company or land manager needs. Effective wildlife management requires that conservation managers can detect changes in how particular target species use a landscape, how this changes over time, understand what is driving these changes, and act to influence these drivers. Increasingly, such adaptive management is a regulatory requirement to access finance or certain markets. For example, the International Finance Corporation’s Performance Standard 6, which is an important guiding framework in international finance, requires that clients “should adopt a practice of adaptive management in which the implementation of mitigation and management measures are responsive to changing conditions and the results of monitoring throughout the project’s lifecycle” [35]. Similarly, the Roundtable on Sustainable Palm Oil’s Principles and Criteria require that the status of rare, threatened and endangered species is monitored and that the “outcomes of this monitoring are fed back into the management plan” [8]. Despite these requirements, and the many companies adhering on paper to them, the reality is that most companies have not gone beyond using presence rather than likelihood of presence of species to guide their management (Meijaard et al. 2020). Such species lists are poor guides to adaptive management, because the only alternative state of presence is absence. Once absence of a previously present species is determined, adaptive management is too late by definition.

### Caveats and statistical validity

While the PENDAKI method has resulted in a large dataset of wildlife observation and trend analyses, there are several issues complicating the statistical quality of the results. Firstly, forest survey blocks are much larger than non-natural sites, which may lead to spurious results. Secondly, there is no fixed species list that observers always record, which may complicate the generation of non-detections.

Thirdly, the data for a considerable number of species are quite poor because they are rarely detected or frequently misidentified. Fourthly, there is likely to be preferential sampling of particular sites which may affect inferences drawn from the data. This relates to the more frequent use of certain roads or survey paths through the plantations. Plans to overcome these difficulties are underway: (1) recording spatial coordinates of observations using the PENDAKI champion application, making it possible to apply smaller forest subsites; (2) Using fixed species lists by a selected sub-group of particularly active observers (started September 2023); (3) The PENDAKI champions are now collecting more data for undersampled species and sites; (4) further improvements in data management, curation and statistical analysis.

Our analysis of naïve versus occupancy-based species and diversity estimates shows that the naïve diversity estimates that companies often use to indicate the conservation value of their land holdings are strongly influenced by survey effort, whereas occupancy-based estimates are not (note the low value from naïve estimates in Fig. 5A for the year 2024 for which only 6 months of data were available). Using naïve estimates, there appears to be an increase in the number of species, which may indicate that the survey and recording effort has increased over time (Fig. 5). Also, this naïve approach underestimates the number of species per block that likely occurs, because not all species are observed in all blocks. The occupancy-based model compensates for this, explaining why dark green bars in Fig. 4 are higher than the light green bars. Both approaches show an increase over time in species richness, which could indeed reflect an increase in species diversity (for example because growing oil palms become ecologically richer with more epiphytes developing over time), or it could reflect a growing species survey and identification ability of observers. In theory, occupancy models can cope with changes in observer capacity over time. If the experience of observers increases, this is reflected in an increasing detection probability and this is corrected for when estimating occupancy. In practice this only works if sufficient data are available per year to estimate detection. In the early year of the PENDAKI data collection, little or no attention was paid to certain species, because the observers did not properly recognize these species, for example Plantain Squirrel (*Callosciurus notatus*) and Prevost’s Squirrel (*C. prevosti*). As a result, the data in the first few years are too poor to be very useful for some species.

### Implementation capacity

PENDAKI (and the later PENDAKI Champion) program has been mostly implemented through ANJ’s own initiative. RD and EM provided the initial idea, technical training, and independent verification, but the actual development of the program, internal training, reward system, app and database development, and external promotion was done by the company itself. The only component, excepting independent verification, for which the company remains dependent on external input is statistical analysis. A training program is currently being implemented to further internalize these processes. The less dependent a company is on external inputs, the easier it would be to implement and maintain it, or scale it up to other companies.

### Cost effectiveness

Compared to other robust wildlife monitoring methods, citizen science-based systems such as PENDAKI are relatively cheap because they are based on voluntary contributions, and neither require expensive technology nor external technical expertise. The long-term annual running costs of USD 0.14/ha compare favourably to traditional wildlife survey methods. For example, a five-year survey effort of a 60,000 ha property in Australia using traditional methods (specialist surveys, animal traps etc.) resulted in 21,000 individual vertebrate records of 216 species for a total cost of about AUS$ 1 million (ca. USD 900,000 in 2009) [36]. This monitoring investment, equivalent to USD 15/ha, was considered insufficient for detecting species populations trends [36]. An alternative monitoring approach using ca. 10 camera traps over 2 years in an area <50 ha in the Netherlands resulted in 47,597 automatically identified species observations at a cost of EUR 66,170/y [37], equivalent to ca. USD 1,300/ha. The costs of PENDAKI are more in line with reported costs of taxonomic surveys, which provide insight into species diversity but not occupancy or abundance, such as USD 0.19/ha survey expenditures in a 500,000 ha Brazilian forest area [38].

### Next steps

To better understand the statistical models, we aim to conduct several studies testing the assumptions in the modelling and also verifying occupancy outputs with ground-truthed data using more traditional survey methods. That might, for example, include an orangutan survey using thermal drones. Through case studies of particular species, we also want to see how the company can operationalize 24/7 adaptive management, where a continuous process of data collection, storage, verification, and analysis, provides estate managers with the kind of practical management feedback that they can easily incorporate in their annual or shorter time-frame management plans. That way, biodiversity management can become an integral part of estate planning and management in a way that it now rarely is in palm oil companies [39].

## Supporting information

S1 Table. Total number of respondents based on location and gender

S2 Table. Accuracy assessment drop-down options

S3 Table. Number of PENDAKI records per taxonomic class, and most frequently recorded taxonomic families

## References

1. Descals A, Gaveau DLA, Serge W, Szantoi Z, Meijaard E. Global mapping of oil palm plantation age for 2021. Earth System Science Data. 2024. doi: 10.5194/essd-16-5111-2024

2. Fitzherbert EB, Struebig MJ, Morel A, Danielsen F, Brulh CA, Donald PF, et al. How will oil palm expansion affect biodiversity? Trends in Ecology & Evolution. 2008;23(10):538–45. doi: 10.1016/j.tree.2008.06.012. PubMed PMID: 2008-361RM-0005

3. Meijaard E, Garcia-Ulloa J, Sheil D, Carlson K, Wich SA, Juffe-Bignoli D, et al., editors. Oil Palm and Biodiversity. A situation analysis by the IUCN Oil Palm Task Force. Gland, Switzerland: IUCN Oil Palm Task Force; 2018. doi: 10.2305/IUCN.CH.2018.11.en

4. Scriven SA, Carlson KM, Hodgson JA, McClean CJ, Heilmayr R, Lucey JM, et al. The Impact of RSPO Membership on Avoiding Biodiversity Losses in Oil Palm Landscapes. Socially and Environmentally Sustainable Oil Palm Research (SEnSOR) Programme, 2017.

5. Lee JSH, Miteva DA, Carlson KM, Heilmayr R, Saif O. Does oil palm certification create trade-offs between environment and development in Indonesia? Environmental Research Letters. 2020;15(12):124064. doi: 10.1088/1748-9326/abc279

6. Meijaard E, Brooks TM, Carlson KM, Slade EM, Garcia-Ulloa J, Gaveau DLA, et al. The environmental impacts of palm oil in context. Nature Plants. 2020;6(12):1418–26. doi: 10.1038/s41477-020-00813-w

7. HCV Resource Network. What are High Conservation Values? https://www.hcvnetwork.org/about-hcvf. 2017.

8. RSPO. RSPO Principles & Criteria Certification For the Production of Sustainable Palm Oil. 2018. Kualu Lumpur, Malaysia: Roundtable on Sustainable Palm Oil, 2018.

9. Furumo PR, Barrera-Gonzalez EI, Espinosa JC, Gómez-Zuluaga GA, Aide TM. Improve Long-Term Biodiversity Management and Monitoring on Certified Oil Palm Plantations in Colombia by Centralizing Efforts at the Sector Level. Frontiers in Forests and Global Change. 2019;2. doi: 10.3389/ffgc.2019.00046

10. Silvy NJ. The Wildlife Techniques Manual. Volume 2 “Management”, 8th Edition. Baltimore John Hopkins University Press; 2020.

11. Silvertown J. A new dawn for citizen science. Trends in Ecology & Evolution. 2009;24(9):467–71. doi: 10.1016/j.tree.2009.03.017

12. Bonney R, Cooper CB, Dickinson J, Kelling S, Phillips T, Rosenberg KV, et al. Citizen Science: A Developing Tool for Expanding Science Knowledge and Scientific Literacy. BioScience. 2009;59(11):977–84. doi: 10.1525/bio.2009.59.11.9

13. Sauermann H, Vohland K, Antoniou V, Balázs B, Göbel C, Karatzas K, et al. Citizen science and sustainability transitions. Research Policy. 2020;49(5):103978. doi: 10.1016/j.respol.2020.103978.

14. Bonney R, Shirk JL, Phillips TB, Wiggins A, Ballard HL, Miller-Rushing AJ, et al. Next Steps for Citizen Science. Science. 2014;343(6178):1436. doi: 10.1126/science.1251554

15. Chandler M, See L, Copas K, Bonde AMZ, López BC, Danielsen F, et al. Contribution of citizen science towards international biodiversity monitoring. Biological Conservation. 2017;213:280–94. doi: 10.1016/j.biocon.2016.09.004

16. Crall AW, Newman GJ, Stohlgren TJ, Holfelder KA, Graham J, Waller DM. Assessing citizen science data quality: an invasive species case study. Conservation Letters. 2011;4(6):433–42. doi: 10.1111/j.1755-263X.2011.00196.x

17. West S, Pateman R. Recruiting and Retaining Participants in Citizen Science: What Can Be Learned from the Volunteering Literature? Citizen Science: Theory and Practice. 2016 1(2):1–10. doi: 10.5334/cstp.8

18. R Core Development Team. R: A language and environment for statistical computing. R Foundation for Statistical Computing. http://www.R-project.org/. Vienna, Austria: 2022.

19. ESRI. Environmental Systems Research Institute, Inc. 2024.

20. MacKenzie DI, Nichols JD, Royle JA, Pollock KH, Hines JE, Bailey LL. Occupancy estimation and modeling: inferring patterns and dynamics of species occurrence (second edition 2018). San Diego, USA: Elsevier; 2006.

21. van Strien AJ, van Swaay CAM, Termaat T. Opportunistic citizen science data of animal species produce reliable estimates of distribution trends if analysed with occupancy models. Journal of Applied Ecology. 2013;50(6):1450–8. doi: 10.1111/1365-2664.12158

22. Ancrenaz M, Oram F, Ambu L, Lackman I, Ahmad E, Elahan H, et al. Of pongo, palms, and perceptions – A multidisciplinary assessment of orangutans in an oil palm context. Oryx. 2015;49(3):465–72. doi: 10.1017/S0030605313001270

23. Ancrenaz M, Oram F, Nardiyono, Silmi M, Jopony MEM, Voigt M, et al. Importance of orangutans in small fragments for maintaining metapopulation dynamics. Frontiers in Forests and Global Change. 2021;4:560944. doi: 10.3389/ffgc.2021.560944

24. Collen BEN, Loh J, Whitmee S, McRae L, Amin R, Baillie JEM. Monitoring Change in Vertebrate Abundance: the Living Planet Index. Conservation Biology. 2009;23(2):317–27. doi: 10.1111/j.1523-1739.2008.01117.x

25. van Strien AJ, Soldaat LL, Gregory RD. Desirable mathematical properties of indicators for biodiversity change. Ecological Indicators. 2012;14(1):202–8. doi: 10.1016/j.ecolind.2011.07.007.

26. Oram F, Kapar MD, Saharon AR, Elahan H, Segaran P, Poloi S, et al. “Engaging the Enemy”: Orangutan (*Pongo pygmaeus morio*) Conservation in Human Modified Environments in the Kinabatangan floodplain of Sabah, Malaysian Borneo. International Journal of Primatology. 2022;43:1–28. doi: 10.1007/s10764-022-00288-w

27. Clary EG, Snyder M. The Motivations to Volunteer: Theoretical and Practical Considerations. Current Directions in Psychological Science. 1999;8(5):156–9. doi: 10.1111/1467-8721.00037

28. Bell S, Marzano M, Cent J, Kobierska H, Podjed D, Vandzinskaite D, et al. What counts? Volunteers and their organisations in the recording and monitoring of biodiversity. Biodiversity and Conservation. 2008;17(14):3443–54. doi: 10.1007/s10531-008-9357-9

29. Lotfian M, Ingensand J, Brovelli MA. A Framework for Classifying Participant Motivation that Considers the Typology of Citizen Science Projects. ISPRS International Journal of Geo-Information [Internet]. 2020; 9(12). doi: 10.3390/ijgi9120704

30. Sutherland W. Evidence-based Conservation. Conservation in Practice. 2003;4(3):39-42. doi: 10.1111/j.1526-4629.2003.tb00068.x

31. Game E, Meijaard E, Sheil D, McDonald-Madden E. Conservation in a wicked complex world; challenges and solutions. Conservation Letters. 2014;7(3):271–7. doi: 10.1111/conl.12050

32. Meijaard E, Sheil D. The dilemma of green business in tropical forests: how to protect what it cannot identify. Conservation Letters. 2012;5(5):342–8. doi: 10.1111/j.1755-263X.2012.00252.x

33. Danielsen F, Ali N, Andrianandrasana HT, Baquero A, Basilius U, de Araujo Lima Constantino P, et al. Involving citizens in monitoring the Kunming–Montreal Global Biodiversity Framework. Nature Sustainability. 2024. doi: 10.1038/s41893-024-01447-y

34. Zuckerberg B, Porter WF, Corwin K. The consistency and stability of abundance–occupancy relationships in large-scale population dynamics. Journal of Animal Ecology. 2009;78(1):172–81. doi: 10.1111/j.1365-2656.2008.01463.x

35. IFC. Performance Standard 6. Biodiversity Conservation and Sustainable Management of Living Natural Resources. International Finance Corporation (IFC), 2012.

36. Perkins GC, Kutt AS, Vanderduys EP, Perry JJ. Evaluating the costs and sampling adequacy of a vertebrate monitoring program. Australian Zoologist. 2013;36(3):373–80. doi: 10.7882/AZ.2013.003

37. Kissling WD, Evans JC, Zilber R, Breeze TD, Shinneman S, Schneider LC, et al. Development of a cost-efficient automated wildlife camera network in a European Natura 2000 site. Basic and Applied Ecology. 2024;79:141–52. doi: 10.1016/j.baae.2024.06.006

38. Gardner TA, Barlow J, Araujo IS, Ávila-Pires TC, Bonaldo AB, Costa JE, et al. The cost- effectiveness of biodiversity surveys in tropical forests. Ecology Letters. 2008;11(2):139–50. doi: 10.1111/j.1461-0248.2007.01133.x.

39. Meijaard E, Ancrenaz M, van Balen S. Biodiversity impact of RSPO certification - an assessment of good practices. Kuala Lumpur, Malaysia: RSPO, 2020.

